# A comprehensive single-cell breast tumor atlas defines cancer epithelial and immune cell heterogeneity and interactions predicting anti-PD-1 therapy response

**DOI:** 10.1101/2022.08.01.501918

**Authors:** Lily Xu, Kaitlyn Saunders, Hildur Knutsdottir, Kenian Chen, Julia Maués, Christine Hodgdon, Evanthia T. Roussos Torres, Sangeetha M. Reddy, Lin Xu, Isaac S. Chan

## Abstract

We present an integrated single-cell RNA-seq resource of the breast tumor microenvironment consisting of 236,363 cells from 119 biopsy samples across 8 publicly available datasets. In this computational study, we first leverage this novel resource to define cancer epithelial cell heterogeneity based on two clinically relevant markers and identify six new and distinct subsets of natural killer cells. We then illustrate how cancer epithelial cell heterogeneity impacts immune cell interactions. We develop T cell InteractPrint, which considers how cancer epithelial cell heterogeneity shifts the predicted strength of T cell interactions. We use InteractPrint to predict response to immune checkpoint inhibition (ICI) in two clinical trials testing immunotherapy in patients with breast cancer. T cell InteractPrint was predictive in both trials (AUC = 0.81 and 0.84), versus PD-L1 expression (AUC = 0.54 and 0.72). This result provides an alternative predictive biomarker to PD-L1 to select patients who should receive ICI.

**STATEMENT OF SIGNIFICANCE:** We developed a novel integrated single-cell atlas of the breast tumor microenvironment to interrogate breast tumor cell heterogeneity and define how heterogenous cancer epithelial cell and immune cell interactions predict response to anti-PD-1 therapy.

## INTRODUCTION

Breast cancer is the most common cancer among women (1). The development of breast cancer is driven by both cancer epithelial cell intrinsic factors and the tumor microenvironment. The medical treatment of breast cancer therefore targets these diverse cell populations and includes traditional chemotherapy, targeted agents inhibiting cancer cell hormone receptors, kinases, cell cycle entry, and immune cell modulators. To further improve these therapies, a deeper understanding of the cellular and molecular composition of breast tumors is required.

Single-cell RNA sequencing (scRNA-seq) technology has been applied to better characterize tumor microenvironments. In breast cancer, several scRNA-seq studies have been performed to identify key immune, cancer cell, and stromal populations of the breast tumor microenvironment (2–9). These studies individually provided insight into the molecular phenotypes of cancer cells, multiple immune populations, and other stromal cells. However, each study was limited by the number of samples and cells analyzed, preventing a comprehensive analysis of rare cell populations as well as the implications of cancer epithelial cell heterogeneity across a larger and more diverse set of clinical breast cancer subtypes.

For example, natural killer (NK) cells are innate lymphoid immune cells critical to anti-tumor defense. In breast cancer, they often represent 1-6% of total tumor cells. Their cytotoxic activity is regulated by a series of functionally activating and inactivating receptors. After tumor exposure, the balance of activating and inactivating receptors can change, and they can lose their cytotoxic activity, proliferative capacities, or even become tumor-promoting (10–12). Because of the small numbers of NK cells processed in most human studies, scRNA-seq analyses of NK cells often are underpowered to capture these distinct functional phenotypes.

In this computational study, we created an integrated atlas of 8 publicly available scRNA-seq datasets from samples taken from patients with early-stage breast cancer (2–9). This resource enables unbiased separation of distinct cell populations within primary breast tumors and robust characterization of phenotypes at the cell level. We then used this resource to define both cancer epithelial cell heterogeneity and rare immune cell heterogeneity along with their subsequent interactions. This comprehensive dataset of the breast tumor microenvironment provides a resource to understand the composition of breast cancer. It is the first to our knowledge to categorize NK cells in breast cancer at the single-cell level and provide evidence that cancer epithelial cell heterogeneity influences immune interactions and response to anti-PD-1 therapy.

## RESULTS

### An integrated scRNA-seq dataset of breast cancer samples reveals the heterogeneity of clinically actionable targets across patients

To develop an atlas of the breast tumor microenvironment that would be statistically powered to capture rare cell populations, we obtained scRNA-seq data from 119 samples collected from primary tumor biopsies of 88 patients across 8 publicly available breast cancer datasets (Fig. 1A; Supplemental Figs. 1A-1C; Supplemental Table 1) (2–9). After processing each dataset separately to filter out low quality cells and doublets, we integrated a total of 236,363 cells across all breast cancer clinical subtypes and a wide spectrum of clinical features (Supplemental Table 1). We assessed batch effect by examining output plots from principal component analysis (PCA) to ensure no cluster was driven by a single dataset or technology (Supplemental Figs. 1D-1E). Cell types were identified by taking the top call resulting from a three-step process which labeled clusters based on a signature score of canonical cell markers, marker count coupled with average expression, and greatest average expression of the marker genes alone (Supplemental Table 2; see Methods). The uniform manifold approximation and projection (UMAP) visualization showed clustering of cells by lineage. Immune cells clustered together across different subtypes while cancer epithelial cells clustered by clinical subtype (Fig. 1B; Supplemental Fig. 1F), which is consistent with other datasets (6, 8).

**Figure 1.**
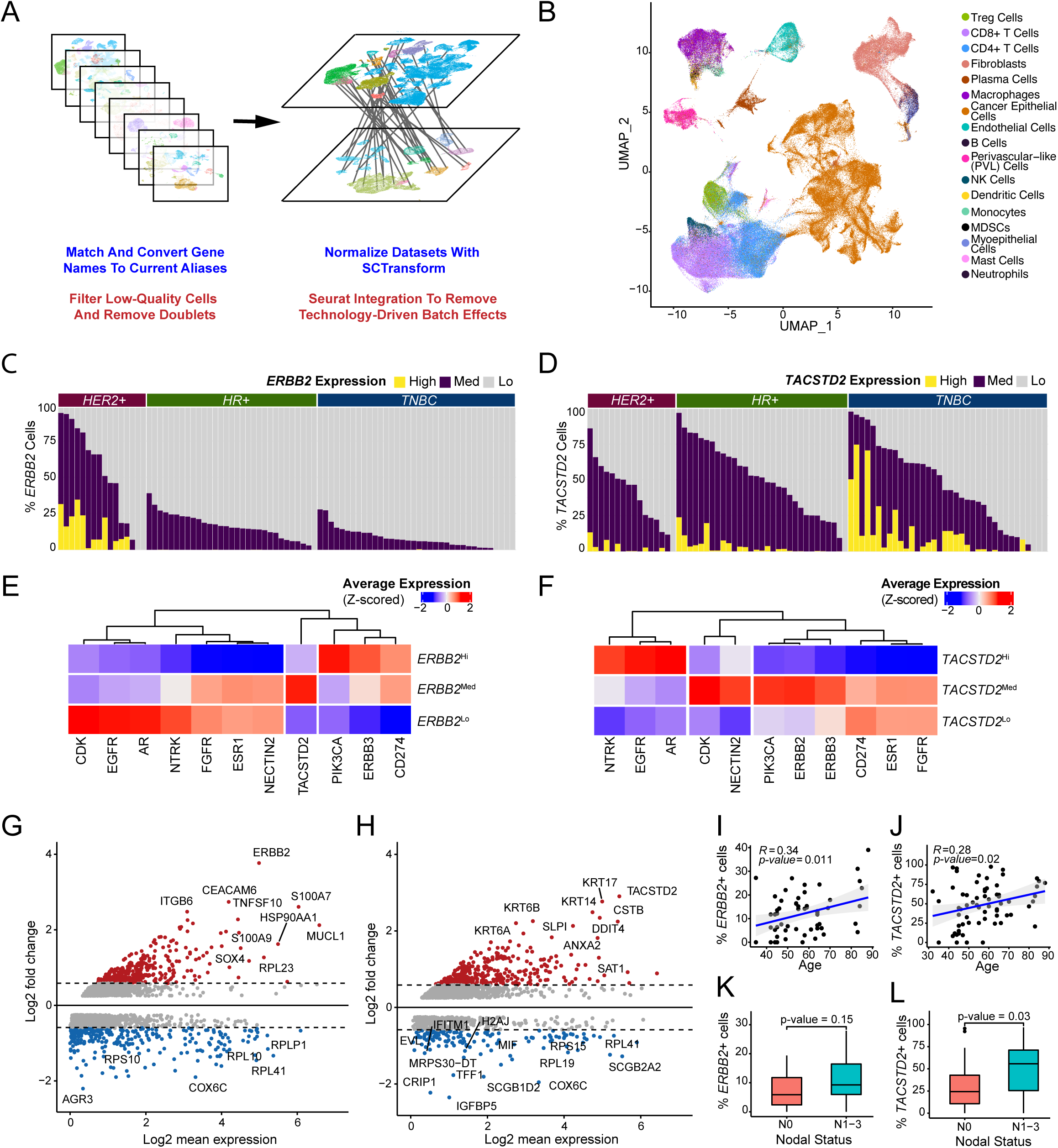
Integrated scRNA-seq dataset of primary breast cancer identifies heterogeneous cancer epithelial cells expressing clinically relevant targets. A. Brief overview of processing and integration pipeline for 8 primary breast cancer datasets. B. UMAP visualization of 236,363 cells across 119 samples from 88 patients analyzed by scRNA-seq. C. Bar plot showing proportion of *ERBB2*^Hi^, *ERBB2*^Med^, and *ERBB2*^Lo^ cells by sample. D. Bar plot showing proportion of *TACSTD2*^Hi^, *TACSTD2*^Med^, and *TACSTD2*^Lo^ cells by sample. E. Heatmap of z-scored average expression of clinically actionable targets in *ERBB2*^Hi^, *ERBB2*^Med^, *ERBB2*^Lo^ cells. F. Heatmap of z-scored average expression of clinically actionable targets in *TACSTD2*^Hi^, *TACSTD2*^Med^, and *TACSTD2*^Lo^ cells. G. MA plot showing differentially expressed genes between *ERBB2*^Hi^ vs. *ERBB2*^Med^ and *ERBB2*^Lo^ cells (Bonferroni adjusted p-value < 0.05). H. MA plot showing differentially expressed genes between *TACSTD2*^Hi^ vs. *TACSTD2*^Med^ and *TACSTD2*^Lo^ cells (Bonferroni adjusted p-value < 0.05). I. Scatterplot showing the Pearson correlation of age and the proportion of *ERBB2*-expressing cells by sample (p-value < 0.05). J. Scatterplot showing the Pearson correlation of age and the proportion of *TACSTD2*-expressing cells by sample (p-value < 0.05). K. Boxplot showing the proportion of *ERBB2*-expressing cells per sample by nodal status (two-sided Wilcoxon test p-value > 0.05). L. Boxplot showing the proportion of *TACSTD2*-expressing cells per sample by nodal status (two-sided Wilcoxon test p-value < 0.05).

To demonstrate the utility of this computational resource, we leveraged this dataset to define heterogeneity among cancer epithelial cells based on expression of genes related to HER2 (*ERBB2*) and TROP2 (*TACSTD2*). This analysis is clinically meaningful because newer anti-HER2 and anti-TROP2 agents show benefit in patients across clinical breast cancer subtypes (13, 14). This recent shift in therapeutic approach highlights a need to better define HER2 and TROP2 expression at the single-cell level to improve patient selection. In contrast to bulk RNA-seq, which have limited resolution due to averaging expression levels across all cell types, this single-cell dataset can be used define interpatient HER2 and TROP2 heterogeneity at the cellular level. While previous bulk RNA-seq and IHC studies have reported expression of the *ERBB2* gene or HER2 protein in up to 70% of HER2-negative breast tumors (15, 16), we were able to detect *ERBB2* expression in 92% of samples independent of clinical subtype at the single-cell level (Fig. 1C). Using a similar analysis for *TACSTD2*, we again found notable heterogeneity (Fig. 1D). In particular, *TACSTD2* expression was observed in 94% of samples across all subtypes, providing additional evidence at single-cell resolution to what has been previously described (17, 18). Moreover, the proportion of *ERBB2*^Hi^ and *ERBB2*^Med^ cells and *TACSTD2^Hi^* and *TACSTD2*^Med^ cells varied between samples. We next asked how other clinically relevant target genes were related to *ERBB2* expression. We found that *PIK3CA, ERBB3*, and *CD274* expression were highest in *ERBB2*^Hi^ cells (Fig. 1E). In contrast, *TACSTD2* expression was highest in *ERBB2*^Med^ cells and notably lower in *ERBB2*^Hi^ cells. Upon analysis of target genes related to *TACSTD2*, we found *EGFR*, *AR,* and *NTRK* expression were elevated in *TACSTD2*^Hi^ cells (Fig. 1F). *ERBB2, ERBB3, PIK3CA,* and *CDK* expression were highest in *TACSTD2*^Med^ cells and lowest in *TACSTD2*^Hi^ cells. We also observed that *TACSTD2*^Med^ cells highly express *NECTIN2*, a ligand related to *TIGIT*, which hints at potential synergy with anti-TROP2 therapeutics and immune checkpoint inhibition.

We then performed differential gene expression analyses on the *ERBB2*^Hi^, *ERBB2*^Med^, and *ERBB2*^Lo^ populations (Fig. 1G; Supplemental Figs. 2A-2B). Of the upregulated genes that were differentially expressed by *ERBB2*^Hi^ cells, 66 genes have been shown to be direct interactors with *ERBB2* (Supplemental Table 3). For instance, *MUCL1* and *ITGB6* were found to be upregulated in *ERBB2*^Hi^ cells and their associations with proliferation of HER2+ breast cancers have been previously characterized (19, 20). For *TACSTD2*^Hi^, *TACSTD2*^Med^, and *TACSTD2*^Lo^ cells, we performed gene expression analyses (Fig. 1H; Supplemental Figs. 2C-2D; Supplemental Table 4). This identified *KRT14* and *KRT17,* which have been implicated as markers for highly metastatic breast cancer cells (21), as significantly upregulated genes in *TACSTD2*^Hi^ cells. We performed gene set enrichment analysis for the *ERBB2* and *TACSTD2* groups to further characterize function (Supplemental Figs. 2E-2F). Interestingly, when assessing for correlation with clinical features, *ERBB2*- and *TACSTD2*-enriched tumors were positively correlated with age (R = 0.34, p = 0.011; R = 0.28, p = 0.02) (Figs. 1I-1J). *ERBB2*-enriched tumors trended towards significant association with higher nodal status (Fig. 1K), while *TACSTD2-*enriched tumors were significantly associated with higher nodal status (p = 0.03) (Fig. 1L).

### Distinct natural killer cell subsets are identified and exhibit diverse functional characteristics

We next sought to leverage our comprehensive dataset to better characterize rarer immune cell populations in human breast cancer. While natural killer (NK) cells are key mediators of anti-tumor control, our understanding of their varied phenotype and function in the breast tumor microenvironment is limited and incomplete. To address this gap, we re-clustered NK cells from the integrated dataset. Unsupervised graph-based clustering of the NK cell scRNA-seq data uncovered 6 distinct clusters of NK cells (Fig. 2A; Supplemental Fig. 3A-3B).

**Figure 2.**
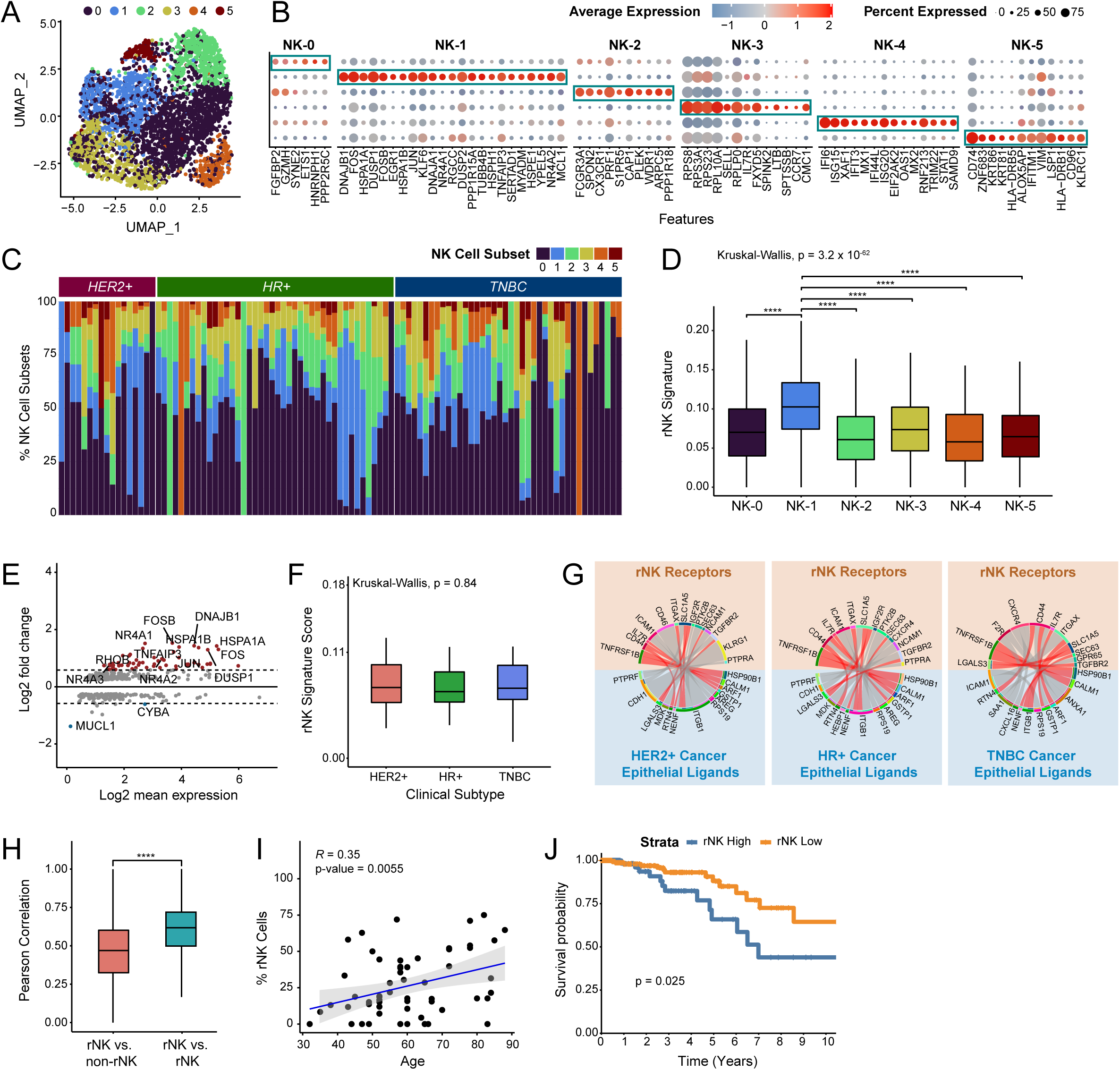
Characterization of natural killer cell subsets in breast cancer. A. UMAP visualization showing major subsets of natural killer (NK) cells. B. Bubble heatmap showing expression of upregulated differentially expressed genes for each major NK cell subset (Bonferroni adjusted p value < 0.05). C. Bar plot showing relative proportions of NK subsets across tumor samples and clinical subtypes. D. Boxplot showing expression of the rNK cell signature in each NK cell subset. NK-1 was significantly different from all other clusters (Kruskal-Wallis p < 0.0001, with post-hoc Dunn test p-values shown; ****p-value < 0.0001). E. MA plot of differentially expressed genes between rNK and non-rNK cells (Bonferroni adjusted p-value < 0.05). F. Boxplot showing the expression level of the rNK signature by clinical subtype. No significant difference was found between subtypes (Kruskal-Wallis p > 0.05). G. Circos plots showing representative predictive receptor-ligand pairs between rNK cells and all cancer epithelial cells separated by clinical subtype. Shared receptors across all subtypes are colored in red. H. Boxplot showing the Pearson correlations of rNK signature gene expression in reprogrammed NK (rNK) cells compared to non-rNK cells versus rNK cells compared to rNK cells (across all clinical subtypes of breast cancer). Pearson correlations between rNK cells and rNK cells are higher than those between rNK cells and non-rNK cells (two-sided Wilcoxon test, ****p-value < 0.0001). I. Scatterplot showing the Pearson correlation of age and proportion of rNK cells by sample (p-value < 0.01). J. Kaplan-Meier plot showing worse clinical outcome in breast cancer patients with high expression of the rNK cell gene signature (log-rank test, p-value < 0.05).

Differential gene expression analysis between clusters revealed significantly upregulated genes defining each NK subset, designated NK-0 through NK-5 (Fig. 2B; Supplemental Figs. 3C; Supplemental Tables 5-6; see Methods). The NK-0, NK-2, and NK-4 populations express genes involved in NK cell cytotoxicity. NK-0 is defined by cytotoxicity-related genes (*GZMH, ETS-1*) (22). NK-2 is defined by genes related to soluble effector functions such as *PRF1* (23) and *CX3CR1* (24). NK-4 is predominated by genes involved in interferon signaling (*IFI6*, *ISG15, XAF1, IFIT3, MX1, IFI144L, ISG20, EIF2AK2, MX2, STAT1*), suggesting that this subset could be involved in the earliest stages of NK cell response to tumor growth (25). NK-3 cells appear to have features of tissue-resident NK cells, having upregulated expression of *SELL* (26), *IL7R* (26), and *CMC1* (27). In contrast, genes of inactivity and reduced cytotoxicity were upregulated in clusters NK-1 and NK-5. NK-1 most notably was marked by genes related to the *NR4A* family (28) and immunosuppressive *DUSP1* (29). Cluster NK-5 expressed genes related to potent inactivating signals of NK cell activity and function, *KLRC1* (30) and *CD96* (31). To further define the function of NK cell subsets, we performed gene set enrichment analysis of individual clusters, which confirmed their functional phenotypes (Supplemental Fig. 3D).

As this was the first time NK cell subsets were described in the breast tumor microenvironment, we sought to define the proportions of subsets within each individual tumor. Overall, we saw a remarkable amount of heterogeneity of NK cell subset proportions, however not all subsets were found within an individual tumor sample (Fig. 2C).

### Reprogrammed NK cells are observed in patient samples independent of subtype

Previously in *ex vivo* and mouse models, we observed that NK cells can be ‘reprogrammed’ after exposure to malignant mammary epithelial cells to promote tumor outgrowth (10, 11). To determine the human significance of this finding, we first generated a signature of mouse reprogrammed NK (rNK) cells based on an experiment (10) comparing the transcriptomes of healthy NK cells to tumor-exposed NK cells that we found to be tumor promoting (Supplemental Fig. 3E). We next converted the original signature to their human analogs (Supplemental Fig. 3F; Supplemental Table 7) and applied it to the NK cell subsets. NK-1 was found to be significantly different from every other NK cell subset (p < 0.0001) (Fig. 2D). Differential gene expression analysis of rNK cells compared to every other NK cell again revealed that the *NR4A* family (*NR4A1*, *NR4A2*, *NR4A3*) were among the most differentially expressed genes (Fig. 2E; Supplemental Table 8; see Methods). *NR4A* are a family of orphan nuclear receptors which act as transcription factors; they are thought to negatively regulate T cell cytotoxicity (32) and have been described as marking specific NK cells that have reduced interferon gamma production (28, 33).

Because of the widespread heterogeneity of NK cell subsets among patients and clinical subtypes, we questioned whether rNK cells were expressed across all breast cancer subtypes. We found no significant differences in rNK cell expression across all clinical subtypes (p > 0.05) (Fig. 2F). Additionally, we found shared receptors across all subtypes (Fig. 2G), which included receptor-ligand pairs *TGFBR2_LGALS3, IL7R_GSTP1,* and *TNFRSF1B_HSP90B1*. Further, we found that the average Pearson correlation in gene expression levels between rNK cells was greater than between rNK and non-rNK cells (p < 0.0001) (Fig. 2H). Together, these findings demonstrate that rNK cells are not defined by specific breast cancer subtype biology, but rather suggest a shared, but still unknown mechanism, contributes to NK cell reprogramming.

To further investigate the clinical significance of rNK cells, we found that higher expression of rNK cells correlates with older age (R = 0.35, p < 0.01) (Fig. 2I). Survival analysis on the TCGA breast cancer cohort showed that increased expression of the rNK cell signature correlates with worse overall survival (Fig. 2J).

### Individual breast tumors have varying degrees of cancer epithelial cell heterogeneity

Given the degree of heterogeneity we observed with single genes and cell types across tumor samples, we sought to understand the degree of cancer epithelial cell heterogeneity within each individual sample. We leveraged the integrated dataset to evaluate the intra-tumoral transcriptional heterogeneity (ITTH) of breast cancer molecular subtypes in an individual patient. Using a well-characterized subtype classifier (8) which scores the four molecular subtypes (Luminal A, Luminal B, Her2, Basal), we found that each patient tumor expressed a varied degree of cells from each molecular subtype (Fig. 3A).

**Figure 3.**
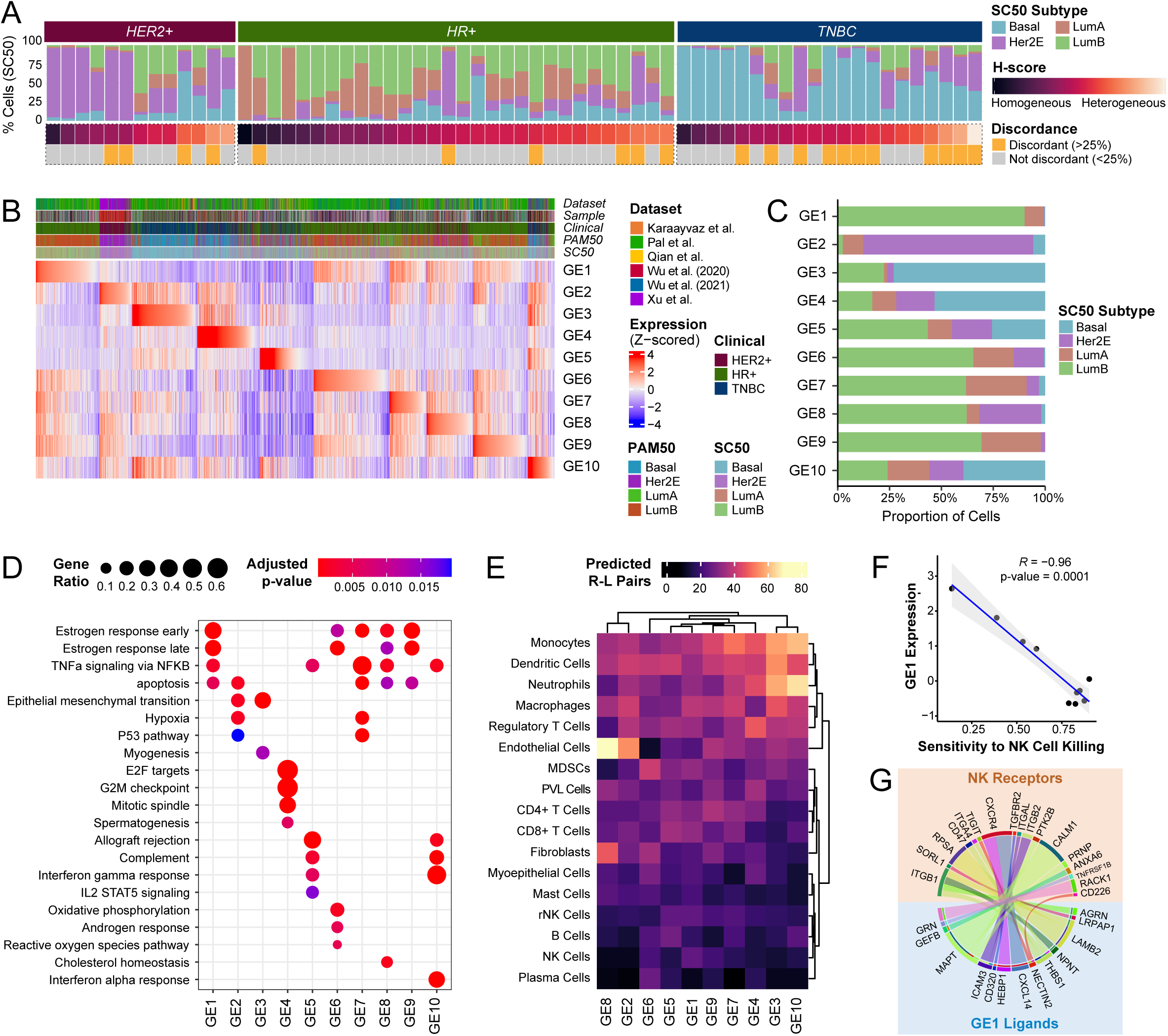
Characterizing the impact of cancer epithelial cell heterogeneity on immune and stromal interactions. A. Percentage of cancer epithelial cells by molecular subtype, sorted by the H-score. Samples with >25% discordance in predicted heterogeneity by molecular subtype and by H-score are denoted. B. Heatmap of z-scored signature scores of the 10 identified gene elements (GEs) representing all cancer epithelial cells, ordered based on the maximum z-scored GE signature score. Annotations represent dataset origin, clinical subtype, PAM50 subtype, and SC50 subtype. C. Percentage of cancer epithelial cells assigned to each GE by molecular subtype. D. Gene set enrichment using ClusterProfiler of the differentially expressed genes by GE. Significantly enriched gene sets from the MSigDB HALLMARK collection are shown (Benjamini-Hochberg adjusted p-value < 0.05). E. Heatmap of prioritized predicted receptor-ligand pairs between cancer epithelial cells by GE and interacting immune and stromal cells. F. Scatterplots showing Pearson correlations of expression of NK-cell related GE1 and sensitivity to NK cell killing (Benjamini-Hochberg adjusted p-value < 0.0005). G. Circos plots showing prioritized receptor-ligand pairs between cancer epithelial cells that highly express NK cell-related GE1 and NK cells.

To better define this phenomenon, we quantified the degree of heterogeneity in expression of non-housekeeping genes across all cancer epithelial cells in a patient tumor, which we termed an ‘H-score’ (Fig 3A). When we applied our H-score to each individual tumor sample, we found that it appeared to visually reflect the degree of molecular subtype heterogeneity. However, we noticed discordance (32.3%) between select patient tumors which demonstrated homogeneity in molecular subtype but high H-scores (see Methods). This suggests that other factors beyond molecular subtype-associated genes drive the observed heterogeneity and underscores a need for different approaches to decode cancer epithelial cell ITTH at higher resolution than that of existing subtype classifiers.

### Cancer epithelial cell heterogeneity can be defined by 10 unifying groups

To develop a high-resolution classifier of heterogeneous cancer epithelial cells, we first performed unsupervised clustering on all cancer epithelial cells in the integrated dataset to generate signatures of upregulated genes that capture distinct molecular features of cancer epithelial cell clusters. Next, supervised clustering was performed based on expression of 12 clinical therapeutic targets (*ESR1*, *ERBB2*, *ERBB3*, *PIK3CA*, *NTRK1/NTRK2/NTRK3*, *CD274*, *EGFR*, *FGFR1/FGFR2/FGFR3/FGFR4, TACSTD2*, *CDK4/CDK6*, *AR*, *NECTIN2*) to ensure that clinically relevant features were captured by upregulated gene signatures and had associations with increased or decreased clinical target expression (see Methods). We additionally supervised clustering of all cancer epithelial cells based on molecular subtype to generate upregulated gene signatures that reflect subtype features. Consensus clustering of all generated gene signatures identified 10 unifying groups, which we defined as ‘gene elements’ (GEs) (Supplemental Fig. 4A). We defined each GE by the top 100 genes that occurred most frequently across gene signatures assigned to the group (Supplemental Table 9; see Methods). We scored each cancer epithelial cell by the individual 10 GEs and assigned GE-based cell labels (Fig. 3B; see Methods). When assessing for molecular subtypes, GE3-labeled cells were predominantly assigned to the basal subtype, while GE2-labeled cells were mainly assigned to the Her2 subtype (Fig. 3C). Cells labeled by GE1 and GE9 were almost exclusively assigned as luminal A and luminal B. In contrast, GE5- and GE10-labeled cells were assigned to all molecular subtypes. Next, we used gene set enrichment analysis to identify functional annotations for each GE (Fig. 3D). GE4 was uniquely enriched for hallmarks of cell cycle and proliferation (*MKI67, PCNA, CDK1*), while GE2 and GE3 contained hallmark genes of EMT (*VIM, ACTA2*). GE1, GE6, GE7, GE8, and GE9 contained genes associated with estrogen response (*ESR1, AREG, TFF3*). GE5 and GE10 were enriched for hallmarks of interferon response (*CD74, ISG15*), allograft rejection (*HLA-DMA, HLA-DRA*), and complement (*C1QA/B/C, C1R*).

To test our hypothesis that GE-based cell labels allow us to characterize cancer epithelial cell heterogeneity within a tumor sample at higher resolution than existing classifiers, we applied our GEs to the integrated dataset to deconstruct each individual patient tumor into the 10 GEs (Supplemental Fig. 4B). Notably, GE-based heterogeneity was not constrained by clinical or molecular subtype. Overall, we generated 10 GEs to characterize cancer epithelial cell ITTH and deconstruct a heterogeneous tumor into its diverse cellular phenotypes.

### Gene elements predict individual patient predominant immune response

We then reasoned that cancer epithelial cell ITTH would also impact each immune interaction within the tumor, where each cancer epithelial cell and immune cell interaction should also be subject to similar degrees of heterogeneity. To examine how cancer epithelial cell ITTH influences immune interactions in the tumor microenvironment, we generated a reference matrix of predicted GE-immune interaction strength by determining the number of predicted receptor-ligand pairings between GEs and immune cells (Fig. 3E; see Methods).

To validate our GE-immune interaction reference matrix, we sought to test these predictions on a separate dataset consisting of human primary breast cancer epithelial cell lines. Cancer epithelial cells labeled by GE1 were predicted to highly interact with NK cells given (highest number of prioritized receptor-ligand pairings). We first applied our GEs to the human primary breast cancer epithelial cell lines to identify which cell lines highly express each GE (Supplemental Fig. 4C). Using a paper by Sheffer et. al. which tests the susceptibility of various cell lines to NK cell cytotoxicity, we assessed the relationship between GE expression and functional cell line sensitivity to NK cell killing (34, 35). As expected, increased GE1 expression was significant correlated with increased resistance to NK cell killing (R = −0.96, p < 0.001) (Fig. 3F). GEs with fewer predicted NK cell interactions did not have statistically meaningful correlations with sensitivity to NK cells (Supplemental Fig. 4D). To investigate reasons underlying these phenotypes, we assessed the predicted receptor-ligand pairing between cells that highly express GE and NK cells (Fig. 3G). We observed that GE1 -labeled cells were predicted to have receptor-ligand pairs that have been characterized as inactivators of NK cell activity (*TIGIT_NECTIN2*, *CD47_THBS1*). This functional study demonstrates that the GEs and the GE-immune interaction reference matrix are predictive of significant NK cell activity with breast cancer cell lines. Overall, this GE-immune interaction reference matrix provides a blueprint for quantifying the degree of interactions between each GE and different immune cell types. Moreover, our receptor-ligand predictions reveal key activating and inhibitory receptors that can be used to infer how GE-immune interactions affect immune cell behavior.

### Spatial mapping of gene elements reflects predicted immune interactions

To validate whether other immune cell interactions could be predicted using our GE-immune interaction reference matrix, we applied our GEs to a spatial transcriptomics dataset published by Wu et al. (8). We first deconvoluted the underlying composition of cell types through integration of the spatial transcriptome data with the integrated dataset (Supplemental Fig. 5A; see Methods). From our GE-immune interaction reference matrix, we were able to infer which GE-labeled cells interact with T cells and which ones do not. Thus, we hypothesized that these GE-labeled cells and CD8+ T cells would be spatially organized in breast tumors. To test this hypothesis, we examined the co-expression of the GEs and the presence of neighboring CD8+ T cells. Notably, GE5 expression demonstrated positive correlations with CD8+ T cells in all samples (mean R *=* 0.26; all p < 0.0001) (Fig. 4A). In one representative image, we determined the co-localization of CD8+ T cells with GE5 expression (Fig. 4B). For areas with high presence of CD8+ T cells (areas 2 and 4), we observed increased colocalization of the predominant receptor-ligand pair (*PTPRC_CD8A*) (Fig. 4B). As expected, GEs with limited predicted interactions did not consistently co-localize with CD8+ T cells (Supplemental Fig. 5B).

**Figure 4.**
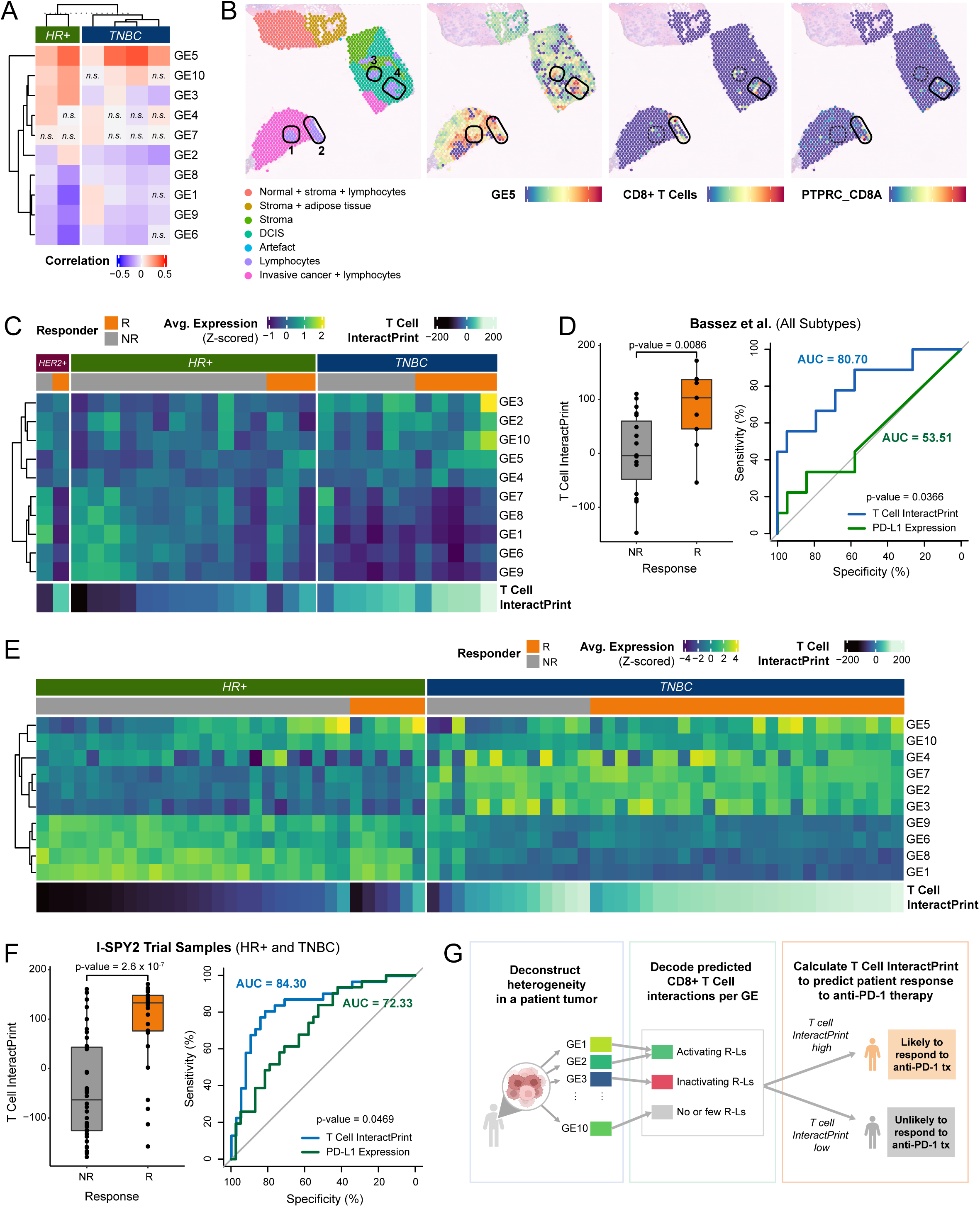
GE-immune interactions predict response to anti-PD-1 therapy. A. Heatmap of Pearson correlations between expression of each of the 10 GEs and the presence of CD8+ T cells for 6 spatial transcriptomics samples across spots containing CD8+ T cells (n.s. Benjamini-Hochberg adjusted p-value > 0.05). B. For a representative TNBC sample, pathological annotation of morphological regions into distinct categories. UCell signature scores of CD8+ T cells overlaid onto spatial tumor sample spots. UCell signature score of GE5 (a CD8+ T cell activating GE) overlaid onto tumor sample spots. Colocalization score for PRPTC and CD8A (a predicted receptor-ligand pair for GE5) overlaid onto tumor sample spots. Tumor sample spots with high predicted presence of CD8+ T cells are outlined. C. Heatmap of average expression of each of the 10 GEs across cancer epithelial cells in each sample from Bassez et al. T cell InteractPrint is shown below. D. Boxplot showing T cell InteractPrint prediction of response to anti-PD-1 therapy across all clinical subtypes in Bassez et al. (R = Responder; NR = Non-responder; p -value < 0.05). AUC of ROC comparing performance of T Cell InteractPrint (AUC = 80.70) and of PD-L1 expression (AUC = 53.5) in Bassez et al. samples (bootstrap test with n = 10,000 p-value < 0.05). E. Heatmap of average expression of each of the 10 GEs across cancer epithelial cells in each sample from the I-SPY2 trial. T cell InteractPrint is shown below. F. Boxplot showing T cell InteractPrint prediction of response to anti-PD-1 therapy across all clinical subtypes in I-SPY 2 trial samples (R = Responder; NR = Non-responder; two-sided Wilcoxon test p-value < 0.0001). AUC of ROC comparing performance of T Cell InteractPrint (AUC = 84.30) and of PD-L1 expression (AUC = 72.33) in I-SPY 2 trial (bootstrap test with n = 10,000 p-value < 0.05). G. Schema of T cell InteractPrint to predict patient response to anti-PD-1 therapy.

### InteractPrint: A weighted score to predict the predominant tumor interacting immune cell for an individual patient tumor

To account for how cancer epithelial cell ITTH within a tumor influences immune cell interactions, we developed InteractPrint. InteractPrint reflects interactions between the predominant tumor-responsive immune cell from Fig. 3E and cancer cells which highly express each GE, weighted by the GE composition of an individual patient tumor. This approach permits real-world application of InteractPrint since it takes into account the heterogeneity of GEs within a tumor.

### InteractPrint predicts anti-PD-1 therapeutic response

We then sought to use InteractPrint to characterize the predominant immune response within patients for therapeutically targeted immune cells. Because current immune checkpoint inhibitors (ICI) target CD8+ T cell-driven cancers, we developed T cell InteractPrint to predict who might respond to ICI.

We applied our approach to a separate scRNA-seq dataset published by Bassez et al., which contains tumor biopsies from breast cancer patients pre- and post-anti-PD-1 therapy (36). Deconstruction of each individual patient tumor into the 10 GEs revealed considerable cancer epithelial cell ITTH prior to anti-PD-1 treatment (Fig. 4C), comparable to levels observed in our integrated dataset (Supplemental Fig. 4B). To assess the capacity of the T cell InteractPrint to predict responders to anti-PD-1 therapy, we derived receiver operating characteristic (ROC) curves in this dataset (Fig. 4D). Across clinical subtypes of breast cancer, the T cell InteractPrint demonstrated an area under the curve (AUC) of 80.70% (p = 0.0086) in predicting response to anti-PD-1 therapy. This was a significant improvement (p = 0.0366) over PD-L1, the current clinical biomarker to predict patients who will respond to anti-PD-1 therapy in breast cancer, which had an AUC of 53.51% (p > 0.05).

Next, we applied our predictor to a separate validation dataset containing results from the I-SPY 2 trial. I-SPY 2 is an ongoing, multicenter, open-label, adaptively randomized phase 2 multicenter trial of neoadjuvant chemotherapy for early-stage breast cancer at high risk of recurrence (37). In this trial, patients with breast cancer received anti-PD-1 therapy (same as patients from Bassez et al.) combined with paclitaxel. We applied our GEs to microarray data from pre-treatment tumor samples from the I-SPY 2 trial and observed levels of GE heterogeneity that were comparable to those described in the scRNA-seq datasets (Fig. 4E). In the I-SPY 2 trial dataset, T cell InteractPrint (AUC = 84.30%; p = 2.6 x 10^-7^) demonstrated significant improvement (p = 0.0469) over PD-L1 (AUC = 72.33%; p = 0.001) in predicting response to anti-PD-1 therapy (Fig. 4F).

Across two trials, T cell InteractPrint demonstrated significant improvement over PD-L1 at predicting response to anti-PD-1 therapy. This highlights the ability of T cell InteractPrint to decode how cancer epithelial cell ITTH impacts CD8+ T cell response for each individual patient.

## DISCUSSION

In our current study, we present a new resource that contains the integration of scRNA-seq data containing 236,363 cells that represent the breast tumor microenvironment. The rate at which scientific data is being generated and published has grown exponentially in the past decade (38). Not only has this led to calls to increase data sharing among research groups (39), but also for more efficient and effective utilization of ‘big data’ to uncover novel biological processes and insights (40, 41). Our approach is to integrate and analyze publicly available data from a single cancer type and apply these analyses to data from functional experiments and clinical trials testing therapies in the same cancer type. This workflow provides a model for how new insights can be derived from utilizing existing data.

We used this resource to improve the characterization of clinically relevant targets in cancer epithelial cells, define subsets of rare immune cell populations that are critical to anti-tumor control, and define how cancer epithelial cell ITTH impacts immune cell interactions across all clinical subtypes of breast cancer. First, we utilize the dataset to identify the heterogenous levels of *ERBB2* and *TACSTD2* expression across all subtypes of breast cancer. This analysis provides additional support for the expansion of therapeutic indication beyond the clinical subtype paradigm. Further, examining genes that are positively correlated with *ERBB2* and *TACSTD2* uncovers other potential clinical targets that can synergize with current anti-HER2 and anti-TROP2 therapies and provides rationale for novel combination approaches.

Then, we leveraged the statistical power of the integrated dataset to identify new subsets of NK cells, consisting of activated and cytotoxic NK cells, exhausted NK cells, and reprogrammed NK cells. Identification of rNK cells in most but not all samples (72%) provides a subtype-independent approach to identify patients who may benefit from rNK cell-directed therapies.

Our results also identify how cancer epithelial cell heterogeneity influences interactions with immune populations. Our schema for T cell InteractPrint allows us to account for, at an individual sample level, how heterogenous expression of GEs shifts the predicted strength of T cell interactions (Fig. 4G). T cell InteractPrint predicted response to T cell immune checkpoint inhibition across all subtypes of breast cancer. This finding is significant, because anti-PD-1 therapy is not effective in HR+ disease (42), and has limited efficacy in TNBC disease (43) when compared to the response seen in other solid tumors (44–47). While the diminished effect of anti-PD-1 in the treatment of breast cancer is multifactorial, one possibility is that the use of PD-L1 is not as sensitive or specific for immune checkpoint inhibition response in breast cancer (5, 48). For example, AUC for PD-L1 in predicting response to anti-PD-1 therapy has been previously estimated to be 65% for breast cancer (49). Even among historically ‘immunogenic’ tumors, PD-L1 AUC is reported to be 70.6% for non-small cell lung cancer and 66.0% for melanoma (50). This result provides a path forward to interpret cancer epithelial cell heterogeneity in a clinically meaningful way.

Future work will be required to further validate our approach to deconstruct cancer epithelial cell heterogeneity and immune interactions. Spatial transcriptomics samples in this study were characterized by limited presence of immune cells, and multiplex IHC or spatial transcriptomics on more clinical samples is required. Application of this same method may allow for predictions of interactions unique to the biology of breast cancer metastasis but will require creation of metastatic sample-specific datasets. While T cell InteractPrint demonstrated significant AUC (p < 0.005) in predicting response to anti-PD-1 therapy, the mechanistic rationale for how cancer epithelial cell ITTH influences CD8+ T cell response to immunotherapy remains unclear, which invites further experimentation.

Our study continues to add to the broader understanding of cancer epithelial cell heterogeneity in the breast tumor microenvironment. We provide further evidence that cancer epithelial cell heterogeneity can be used to predict its role in cancer cell-immune cell interactions.

## METHODS

### Processing of single-cell RNA-Seq Datasets

We obtained 119 primary breast tumor samples across 8 publicly available datasets from 88 untreated female patients ranging from 32 to 90 years of age encompassing all clinical subtypes. As the collected datasets were not aligned to the same genome, all gene names were converted to the official gene alias as delineated by the HUGO Gene Nomenclature Committee (HGNC) using the *limma (v3.50.1)* and *org.Hs.eg.db (v3.14.0)* packages (51, 52). Each dataset was then filtered based on percent mitochondrial transcripts, percent hemoglobin genes, number of RNA molecules, and number of features. In brief, cells lower than the 5^th^ percentile and greater than the 95^th^ percentile of each metric were removed, as well as those cells with greater than 15% mitochondrial content.

To avoid confounding clustering and gene expression analyses, we used the *DoubletFinder (v2.0.3)* package to identify and remove doublets from the dataset (53). Doublet rates were estimated based on given rates from the original technology used and the cell loadings provided by the original studies.

### Integration of Primary Breast Cancer Datasets by Seurat

The 119 untreated primary samples were integrated via reference-based integration using *Seurat (v4.1.0)* to remove any technology-driven batch effects (54). To prevent over-correction of the data, *SCTransform (v0.3.2.9008)* was used rather than the standard Seurat normalization scheme (55). This was done according to the developers’ vignette (https://satijalab.org/seurat/articles/integration_large_datasets.html). The 10x datasets were chosen as the reference and the *rann* method was chosen for *FindNeighbors*. Success of batch effect correction was determined by visually inspecting the top two principal components and ensuring that no single technology was driving any clusters.

### Cell Type Annotation and Clustering

First, general cell types were identified using canonical and literature-derived cell markers as specified in Supplemental Table 2. After these were determined, three methods were used to refine each cell’s identity. The first utilized cluster-level annotations via the *UCell (v1.99.1)* package (56); the second labeled cells based on thresholds based on the number of markers, and then clustered and calculated the average expression of those markers to refine the cell identities (3); and the third took highest average expression of select markers. The annotation with highest agreement across these methods was selected as the cell type. If all methods disagreed, then the overall cluster labeling was chosen as the annotation for that cell.

For the cluster-level method, all cell markers were aggregated into a single score using the *AddModuleScore_UCell* function from the *UCell (v1.99.1)* package and visualized using *FeaturePlot* from the *Seurat (v4.1.0)* package (54, 56). The clusters that had the highest overall UScore for a given cell type were labeled as that type, isolated and re-integrated to account for technology-driven batch effects, and subtype-specific cell markers were applied (e.g., *CD4* for CD4+ T cells).

For the second method, cells were identified as a given type based on the number of markers that had non-zero expression for a given cell. In brief, epithelial cells were labeled as such if they had two epithelial markers or if they had at least one of the following markers: *EPCAM, KRT8, KRT18,* and *KRT19*. Specific immune types were labeled as such if they either had at least two markers of that type and no other type, *PTPRC* and at least one marker of that type and no other, or at least three markers for that type and at most one marker of a different immune type. Stromal cell types could either have only cell-type-specific markers or at least three cell-type-specific markers and at most one endothelial marker. Finally, endothelial cells could have either only endothelial markers or at least three endothelial markers and at most one marker associated with a stromal cell type.

Lastly, we examined log-normalized expression values of the selected markers for each cell. A cell was assigned to the cell type that had the highest average expression for their markers across all features. T and myeloid subsets were identified in the same manner once the cells were identified as T cells or myeloid cells respectively.

The final cell call was determined based on the highest consensus or defaulted to the larger cluster’s identity.

### Identification of Cancer Epithelial Cell Populations

The larger epithelial cluster was re-clustering and re-integrated to account for technology-driven batch effects. Select rare populations (*ERBB2*-positive and *TACSTD2*-positive) were then chosen due to known rarity and clinical relevance. *ERBB2* and *TACSTD2* expression levels are calculated using *UCell (v1.99.1)* (56)*. ERBB2*^Hi^ cancer epithelial cells were defined by positive *ERRB2* expression above the 97.5^th^ percentile, *ERBB2*^Med^ cells were defined by positive *ERBB2* expression at or below the 97.5^th^ percentile, and *ERBB2*^Lo^ cells were defined by zero *ERBB2* expression. *TACSTD2*^Hi^ cells were defined by positive *TACSTD2* expression above the 95^th^ percentile, *TACSTD2*^Med^ cells were defined by positive *TACSTD2* expression at or below the 95^th^ percentile, and *TACSTD2*^Lo^ cells were defined by zero *TACSTD2* expression.

*FindMarkers* in *Seurat (v4.1.0)* and *MAST (v1.20.0)* were then used to identify differentially expressed genes in each cluster (>5 cells per cluster, only test genes detected in >20% of cells in a cluster) (54, 57). Expression levels of clinically actionable targets for each subset of cells was estimated by *AverageExpression* by *Seurat (v4.1.0)* (54). For exploring associations with clinical features, linear regression and Pearson correlations were calculated between the proportion of *ERBB2*-positive or *TACSTD2-*positive cells per sample and the age or nodal status. For visualization of differentially expressed genes, the log_2_ fold change cutoff was increased to 1.5 and a false discovery rate cutoff was set to be equal to 0.05. Gene set enrichment analysis across the *ERBB2* and *TACSTD2* groups was performed using *clusterProfiler (v4.2.2)* and the Hallmark gene set collection from *msigdbr (v7.5.1)* (58, 59) using an additional cutoff of the absolute difference in percent expression between the pairwise populations > 0.1. Only genes with a log_2_ fold change > 0 were considered.

### Identification of Natural Killer Cell Subsets

NK subpopulations were found through the isolation and re-integration of the larger NK cluster gain to ensure the removal of technology-driven batch effects. Given the higher dimensionality of the dataset containing only NK cells, the Manhattan distance metric was used. *FindMarkers* in *Seurat (v4.1.0)* and *MAST (v1.20.0)* were used to identify differentially expressed genes for each cluster (Bonferroni adjusted p value < 0.05) (54, 57). For marker gene selection, the absolute log_2_ fold change cutoff was increased to a baseline of 0.56. Thresholds used to select the highest marker genes for each NK subset are included in Supplemental Table 5.

To identify human reprogrammed tumor-promoting NK cells, we first developed a 99 gene signature that based on genes upregulated in tumor-exposed NK cells as compared to healthy NK cells in MMTV-PyMT and WT FVB/n mice as previously described (10). In that paper, primary healthy and tumor-exposed NK cells were isolated and total RNA was extracted and sequenced using Illumina NextSeq 500. Bulk RNA-seq paired-end reads were then aligned and mapped using hisat2 (60) and HTSeq (61) respectively, and DESeq2 was used for differential gene expression analysis between these two populations.

For application in the current study, the mouse genes were converted into their human aliases using the *BioMart (v2.50.0)* package (62). Because the mouse strain used in the previous study (MMTV-PyMT) most closely resembles the luminal A/luminal B and basal subtypes, these same subtypes were analyzed for rNK presence. NK cells within each subtype that were in the 75^th^ percentile for this 90 gene signature were labeled as “reprogrammed”. Removing any duplicates resulting from HR+ cells being included in the luminal A and luminal B groups led to 841 total rNK cells in the dataset.

Gene set enrichment analysis across the NK subsets was performed using *clusterProfiler (v4.2.2)* and the Hallmark gene set collection from *msigdbr (v7.5.1)* (58, 59). Only genes with a log_2_ fold change > 0 were considered. Samples with fewer than 10 NK cells were omitted from the correlative analyses comparing proportion of rNK cells and clinical variables. For visualization of differentially expressed genes, the log_2_ fold change cutoff was increased to 1.5 and a false discovery rate cutoff was set to be equal to 0.05. To examine expression of the rNK signature within the NK cell subsets and across the clinical subtypes, the Kruskal-Wallis test and a pairwise post-hoc Dunn test was performed when appropriate. For the similarity analysis of rNK cells, the expression matrix was reduced to the reprogrammed signature, and the Pearson correlation coefficient was calculated for all pairwise combinations of rNK cells with rNK cells and for rNK cells with non-rNK cells.

### Molecular Subtyping of Samples Using SCSubtype

To identify the molecular breast cancer subtypes within the integrated dataset, we used the SCSubtype gene signature described in Wu et al. (8).

In brief, the mean read counts for each signature were determined and the highest mean was assigned as the subtype for that cell. To identify the molecular subtype of each sample, we determined the number of cells classified under each subtype, and then picked the subtype with the highest number of cells as the subtype for that sample following the method of Wu et al. (8).

### Gene signature analysis of cancer epithelial cell heterogeneity

For each individual tumor sample with more than 50 epithelial cells, pairwise cell-cell Pearson correlations were calculated across all non-housekeeping genes across all epithelial cells in the sample using *Rfast (v2.0.6)* (63). The ‘H-score’ was calculated as the median of all pairwise Pearson correlations for that sample, normalized across the integrated dataset. To identify discordance between heterogeneity as characterized by the ‘H-score’ vs. by the molecular subtype, we calculated the difference between the ‘H-score’ and the maximum percentage of cells within the sample that belonged to a single subtype. For samples with discordance >25%, molecular subtype classifiers were limited in resolution for defining intratumoral heterogeneity.

Unsupervised and supervised clustering of cancer epithelial cells for each individual tumor sample with more than 50 epithelial cells was performed.

For unsupervised clustering on the integrated dataset, all cancer epithelial cells were clustered at 15 resolutions (0.01, 0.05, 0.08, 0.1, 0.2, 0.3, 0.4, 0.5, 0.6, 0.7, 1.0, 1.3, 1.6, 1.8, 2.0) utilizing *Seurat (v4.1.0)* (54)*. FindMarkers* in *Seurat (v4.1.0)* and *MAST (v1.20.0)* were then used to identify differentially expressed genes in each cluster (>5 cells per cluster, only test genes with >25% difference in the fraction of detection between the clusters, log_2_FC > 0.25) (54, 57). Dataset-wide unsupervised clustering returned 519 differentially expressed gene signatures of upregulated genes.

For unsupervised clustering on the sample level, cancer epithelial cells were clustered by sample at 15 resolutions (0.01, 0.05, 0.08, 0.1, 0.2, 0.3, 0.4, 0.5, 0.6, 0.7, 1.0, 1.3, 1.6, 1.8, 2.0) utilizing *Seurat (v4.1.0)* (54)*. FindMarkers* in *Seurat (v4.1.0)* and *MAST (v1.20.0)* were then used to identify differentially expressed genes in each cluster (>5 cells per cluster, only test genes with >25% difference in the fraction of detection between the clusters, log_2_FC > 0.25) (54, 57). Sample-level unsupervised clustering returned 5,546 differentially expressed gene signatures of upregulated genes.

For supervised clustering by SCSubtype (i.e., basal, HER2-enriched, luminal A, luminal B), epithelial cells were clustered based on SC50 subtype (8). *FindMarkers* in *Seurat (v4.1.0)* and *MAST (v1.20.0)* were then used to identify differentially expressed genes in each cluster (>5 cells per cluster, only test genes detected in >20% of cells in a cluster, log_2_FC > 0.1) (54, 57). Supervised clustering based on clinical target expression returned 4 differentially expressed gene signatures of upregulated genes.

For supervised clustering based on clinical target expression (*ESR1, ERBB2, ERBB3, PIK3CA, NTRK1/NTRK2/NTRK3, CD274, EGFR, FGFR1/FGFR2/FGFR3/FGFR4, TACSTD2, CDK4/CDK6, AR, NECTIN2*), epithelial cells were scored using *UCell (v1.99.1)* and clustered based on high (expression level above the top 90^th^ percentile of all epithelial cells), medium (expression level below the top 90^th^ percentile but greater than zero), and low (no or zero expression level) expression of clinical targets (56). *FindMarkers* in *Seurat (v4.1.0)* and *MAST (v1.20.0)* were then used to identify differentially expressed genes in each cluster (>5 cells per cluster, only test genes detected in >20% of cells in a cluster, log_2_FC > 0.1) (54, 57). Supervised clustering based on clinical target expression returned 32 differentially expressed gene signatures of upregulated genes.

Only gene signatures containing more than 20 genes and originating from clusters of >5 epithelial cells were kept. Additionally, redundancy was reduced by comparing all pairs of unsupervised gene signatures and removing the pair with the fewest genes from pairs with Jaccard index >0.9. Across all tumor samples, a total of 1,101 gene signatures were generated.

Consensus clustering of the Jaccard similarities (using skmeans clustering, ATC, implemented in the *cola (v2.0.0)* package) between these gene signatures was used to identify 10 groupings (64). For each grouping, we took the top 100 genes that have the highest frequency of occurrence across clusters. These were defined as a gene element (i.e., GE) and were named GE1 to GE10. GE signature expression was calculated for each cancer epithelial cell using *UCell (v1.99.1)* (56). For Fig. 3B and 3C, GE signature expression was z-scored across all cancer epithelial cells, and each cell was assigned to the GE with the highest z-scored expression.

Gene set enrichment analysis for each GE was performed using *clusterProfiler (v4.2.2)* and the Hallmark gene set collection from *msigdbr (v7.5.1)* (58,59,65). Only gene sets predicted to be significantly enriched within each GE are shown.

### Survival Analysis

To determine survival outcomes, we obtained the primary solid tumor samples from the breast cancer cohort of The Cancer Genome Atlas (TCGA) Project (66). Genes names were converted using the same method described in the scRNA-seq processing section. The data was normalized using the *TCGAAnalyze_Normalization* function from the *TCGAbiolinks (v2.18.0)* package with default parameters and was subsequently transformed using the *vst* function from the *DESeq2 (v1.34.0)* package, also with default parameters (67, 68).

We assessed survival outcomes on the TCGA samples with high immune infiltrate (activated and resting NK cells predicted to be greater than a relative fraction of 0.015 of tumor-infiltrating immune cells in the sample), as defined by Xu et al. (9). We applied the 44 upregulated genes of the rNK signature to all TCGA samples and labeled the top 300 patients with highest rNK signature expression as ‘rNK-high’ and the bottom 801 patients with lowest rNK signature expression as ‘rNK-low.“ Survival curves were generated using the Kaplan-Meier method with the *survival* package *(v2.44-1.1)* (69). We assessed the significance between two groups using log-rank test statistics.

### Receptor-ligand pairing analysis between cancer epithelial cells and immune cells

To further delineate the commonalities across the subtypes driving NK cell reprogramming, separate *NicheNet* analyses were run between rNK cells and cancer epithelial cells separated by clinical subtype (HER2+, HR+, TNBC), where rNK cells were set as the ‘sender’ population, and non-rNK were set as the ‘reference’) (70). Receptor-ligand regulatory potential scores for the top 50 predicted ligands and top 200 predicted targets were calculated and for each predicted receptor-ligand pair, an R-L interaction score was calculated as a product of ligand expression (fold change in average expression of the ligand in cancer epithelial cells of that clinical subtype) and receptor expression (percent of the rNK population that has positive expression of the receptor). For the top 20 receptor-ligand pairs selected based on this R-L interaction score, circos plots were generated to visualize links between ligands on the cancer epithelial cells by GE and receptors on the interacting cell subsets.

Next, receptor-ligand pairing analysis was performed using *NicheNet (v1.1.0)* and *CellChat (v0.0.1)* to explore interactions between cancer epithelial cells and interacting immune and stromal cells (i.e., CD4+ T cells, CD8+ T cells, regulatory T cells, B cells, plasma cells, myeloid cells, mast cells, MDSCs, NK cells, rNK cells, fibroblasts, myoepithelial cells, endothelial cells, perivascular-like cells) (70, 71).

For each cancer epithelial cell, GE signature expression was z-scored across all cancer epithelial cells, and each cell was assigned to the GE with the highest z-scored expression. For each GE, in a similar manner as with the rNK analyses, multiple separate *NicheNet* analyses were run between cancer epithelial cells assigned to that GE (set as ‘sender’ population) and each interacting cell subset (set as ‘receiver’ population). The top 50 predicted ligands and top 200 predicted targets were used for the development of the R-L interaction score, which was the product of the fold change in average expression of the ligand on cancer epithelial cells with high vs. low GE expression, and receptor expression (percent of the interacting cell subset that has positive expression of the receptor). For the top 20 receptor-ligand pairs selected based on this R-L interaction score, circos plots were again generated to visualize links between ligands on the cancer epithelial cells by GE and receptors on the interacting cell subsets.

In addition to the *NicheNet* analysis, cancer cell and interacting cell communication analysis was conducted using *CellChat (v0.0.1)* with default parameters (71). All cancer epithelial cells were assigned to the GE with the highest z-scored expression. For each GE, the cell-cell communication network between GE-labeled cancer epithelial cells and interacting cells was visualized using *CellChat (v0.0.1)* (104). For each GE and interacting cell pair, receptor-ligand pairings with significant (Bonferroni adjusted p-value < 0.05) probability of interaction are selected.

To infer the degree of interaction between the GEs and various immune or stromal cell populations, we estimate the number of key receptor-ligand interactions for each GE and interacting cell population. First, the list of receptor-ligand interactions predicted by *NicheNet* were filtered. For each interacting cell population, the top 2,000 predicted receptor-ligands for each interacting cell population were selected based on *Nichenet* prediction for regulatory potential. Then, of those selected pairs, the top 400 predicted receptor-ligands for each GE were selected based on ligand expression (fold change in average expression of the ligand on cancer epithelial cells with high vs. low GE expression). Lastly, all overlapping receptor-ligand interactions that were predicted by both *NicheNet* and *CellChat* for a GE and interacting cell pair are selected. We combined the list of overlapping receptor-ligand interactions and the list of selected *NicheNet* receptor-ligand interactions to generate our list of prioritized receptor-ligand interactions for each GE and immune or stromal cell population.

For each GE and interacting cell pair, the number of prioritized receptor-ligand interactions was used to infer the degree of interaction between the GE and the interacting cell population. We capture the number of prioritized receptor-ligand interactions in our GE-immune interaction reference matrix.

### Breast cancer cell line exploration

To explore cancer epithelial cell heterogeneity and NK cell sensitivity, we obtained bulk RNA-seq data for primary breast cancer cell lines from the Broad Cancer Cell Line Encyclopedia (CCLE) differentiated by their sensitivity to NK cells (34, 35). Genes names were converted using the same method described in the scRNA-seq processing section. Normalized expression data were deconvoluted with *BisqueRNA (v1.0.5)* using marker-based devolution with the 10 GE signatures in order to estimate the relative abundance of the GEs for each cell line (72). NK cell sensitivity was assessed using reported 72-hour AUC values for each primary breast cancer cell line. Linear regression and Pearson correlation were used to assess the relationship between GE expression and NK cell sensitivity (34, 35). GE signature expression was calculated for each primary breast cancer cell line (n = 10) using *UCell (v1.99.1)* (56). NK cell sensitivity was assessed using reported 72-hour AUC values for each primary breast cancer cell line. Linear regression and Pearson correlation were used to assess the relationship between GE expression and NK cell sensitivity.

### Spatial transcriptomics analysis

Processed spatial transcriptomics count matrices for 6 samples from Wu et al. were loaded into *Seurat* (v.4.1.0) (8, 54). Because each spot captures multiple cells, we deconvoluted the underlying composition of cell types using the anchor-based Seurat integration workflow. This workflow used *SCTransform* normalization (55) and transferred annotations from the integrated scRNA-seq dataset reference to the spatial transcriptomic datasets. The resulting annotations calculated the fraction of each cell type per given spot and mapped the spatial distribution of cell types, which we further corroborated by the spatial expression of marker genes (Supplementary Table 2). Spatial and scRNA-seq data were matched with respect to breast cancer clinical subtype. Spots labeled as normal tissue or artefact by pathologist annotation were excluded from the analysis.

To investigate interactions between cancer epithelial cells by GE and immune or stromal cells, spots were first filtered based on the predicted scores of the ‘cancer epithelial cell’ annotation called by the *Seurat* integration (spots with less than 10% predicted cancer epithelial cells are excluded) (54). Each spot containing cancer epithelial cells was then scored for expression of each of the 10 GEs using *UCell (v1.99.1)* (*56*). For immune and stromal cell populations, spots were filtered based on predicted scores for their respective annotations called by the *Seurat* integration (spots with 0% predicted cells are excluded) (54). For each immune and stromal cell population, each spot containing the respective cell type was scored for expression of the cell using canonical and literature-derived cell markers as specified in Supplemental Table 2 by the *UCell (v1.99.1)* package (56). To assess GE and CD8+ T cell colocalization for all samples, Pearson correlations were computed across spots containing between the expression of each GE and the expression of CD8+ T cell markers. For cell signaling predictions between select GE ligands and CD8+ T cell receptors, receptor-ligand co-localization scores were defined as the product of the ligand and receptor normalized expression levels.

### Development of InteractPrint, a cancer epithelial cell-immune cell interaction score

For each sample, the average expression of each GE is calculated as the average of the scaled *UCell* score (scaled across all cancer epithelial cells in the dataset). Next, the number of prioritized receptor-ligand interactions in the GE-immune reference matrix between each GE and CD8+ T cells is used to infer the degree of interaction between cancer epithelial cells and CD8+ T cells. GE1, GE6, GE7, GE8, and GE9 were designated as “inactivating” based on the presence inactivating CD8+ T cell receptors (e.g., *NECTIN2_TIGIT*) in the list of prioritized receptor-ligand interactions for those GEs.

For each sample, the T cell InteractPrint was calculated as the average of the number of prioritized CD8+ T cell receptor-ligand interactions in the GE-immune interaction reference matrix, weighted by average expression of each GE and a factor of −1 for inactivating GEs.

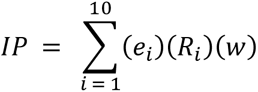

*InteractPrint =* Weighted CD8+ T cell interaction score calculated for a patient tumor

*i =* GE (ranges from 1 to 10)

*e_i_ =* Average GE expression (average of z-scored *UCell* scores for the GE across all cells in the sample)

*R_i_ =* Number of prioritized receptors on the interacting cell type (prioritized from NicheNet and CellChat)

*w =* Multiplier for activating or inactivating GE (*w =* 1 for CD8+ T cell activating GEs; *w = − 1* for CD8+ T cell inactivating GEs)

### Validation of InteractPrint

To assess the predictive value of the T cell InteractPrint, we applied our method to a publicly available scRNA-seq dataset containing 29 primary breast tumors from patients who received pembrolizumab anti-PD-1 therapy (Bassez et al.) (36). GE signature expression was calculated for each pre-treatment sample using *UCell (v1.99.1)* (56), and the T cell InteractPrint was calculated for each sample. We assessed the predictive value of the T cell InteractPrint compared to average expression levels of PD-L1. ROC curves and AUC statistics were generated using the *pROC (v1.18.0)* and default settings (73). Bootstrap method (n = 10,000) in *pROC (v1.18.0)* was used for significance testing between T cell InteractPrint ROC and PD-L1 ROC curves.

To further assess the predictive value of InteractPrint, we applied our method to the I-SPY 2 microarray dataset containing 69 primary breast tumors from patients who received combination paclitaxel and pembrolizumab anti-PD-1 therapy (37). The data was loaded using *limma (v3.15)*, and the batch-corrected and normalized expression data provided by the authors was inserted into the object (51). Genes names were converted using the same method described in the scRNA-seq processing section. Microarray data was deconvoluted with *BisqueRNA (v1.0.5)* using marker-based devolution with the 10 GE signatures in order to estimate the relative abundance of the GEs within each sample (72). GE signature expression was compared for responders and non-responders. We again assessed the predictive value of the T cell InteractPrint compared to scaled expression levels of PD-L1. ROC curves and AUC statistics were generated using the *pROC (v1.18.0)* and default settings (73). The data was loaded using *limma (v3.15)*, and the batch-corrected and normalized expression data provided by the authors was inserted into the object (51). Genes names were converted using the same method described in the scRNA-seq processing section. Microarray data was deconvoluted with *BisqueRNA (v1.0.5)* using marker-based devolution with the 10 GE signatures in order to estimate the relative abundance of the GEs within each sample (72). GE signature expression was compared for responders and non-responders (Supplemental Figure 7). We again assessed the predictive value of the T cell InteractPrint compared to scaled expression levels of PD-L1. ROC curves and AUC statistics were generated using the *pROC (v1.18.0)* and default settings (73).

### Statistical analyses

Statistical significance was determined using the Wilcoxon Rank Sum test (unless otherwise stated in the figure legend). Where appropriate, p-values were adjusted using the Bonferroni correction (unless otherwise stated in the figure legend) where appropriate for multiple testing. All box plots depict the first and third quartiles as the lower and upper bounds, respectively. The whiskers represent 1.5x the interquartile range and the center depicts the median. All statistical tests used are defined in the figure legends. P-values < 0.05 were considered significant.

## Data and Code Availability

Upon publication, all processed scRNA-seq data in the integrated dataset will be made available for in-browser exploration and visualization through the Broad Institute Single Cell portal at https://singlecell.broadinstitute.org/single_cell/. Processed scRNA-seq data from this study will also be made available for download through the Gene Expression Omnibus upon publication.

Six of the nine publicly available scRNA-seq datasets were retrieved from the Gene Expression Omnibus under the following accession codes: GSE114727 (Azizi et al. (2)), GSE118389 (Karaayvaz et al. (3)), GSE161529 (Pal et al. (4)), GSE110686 (Savas et al. (74)), GSE176078 (Wu et al. (8)), and GSE180286 (Xu et al. (9)). The remaining three scRNA-seq datasets can be found at the following links: https://lambrechtslab.sites.vib.be/en/single-cell (Bassez et al. (36)), https://lambrechtslab.sites.vib.be/en/pan-cancer-blueprint-tumour-microenvironment-0 (Qian et al. (6)), and https://singlecell.broadinstitute.org/single_cell/study/SCP1106/stromal-cell-diversity-associated-with-immune-evasion-in-human-triple-negative-breast-cancer (Wu et al. (7)). The spatially resolved data from Wu et al. were retrieved from the Zenodo data repository (https://doi.org/10.5281/zenodo.4739739). The breast cancer TCGA expression data was downloaded using the *TCGAbiolinks* package (67). Breast cancer cell lines were downloaded from the depmap portal at the following link: https://depmap.org/portal/download/. The I-SPY2-990 microarray data was retrieved from the Gene Expression Omnibus under the accession code GSE194040 (37).

Code related to these analyses will be made available at https://github.com/ChanLab-UTSW/BreastCancer_Integrated upon publication. Additional data and methods are available from the corresponding author upon request.

## ACKNOWLEDGEMENTS

We would like to thank Drs. Carlos Arteaga, Suzanne Conzen, and John Minna for reading our manuscript and providing helpful feedback. This study was supported by funding provided by METAvivor, the Peter Carlson Trust, Theresa’s Research Foundation, and the NCI/UTSW Simmons Cancer Center P30 CA142543. Special thanks to all members of the Chan Lab.

