## Supplemental Figures for "A comprehensive single-cell breast tumor atlas defines cancer epithelial and immune cell heterogeneity and interactions predicting anti-PD-1 therapy response"

### Figure S1

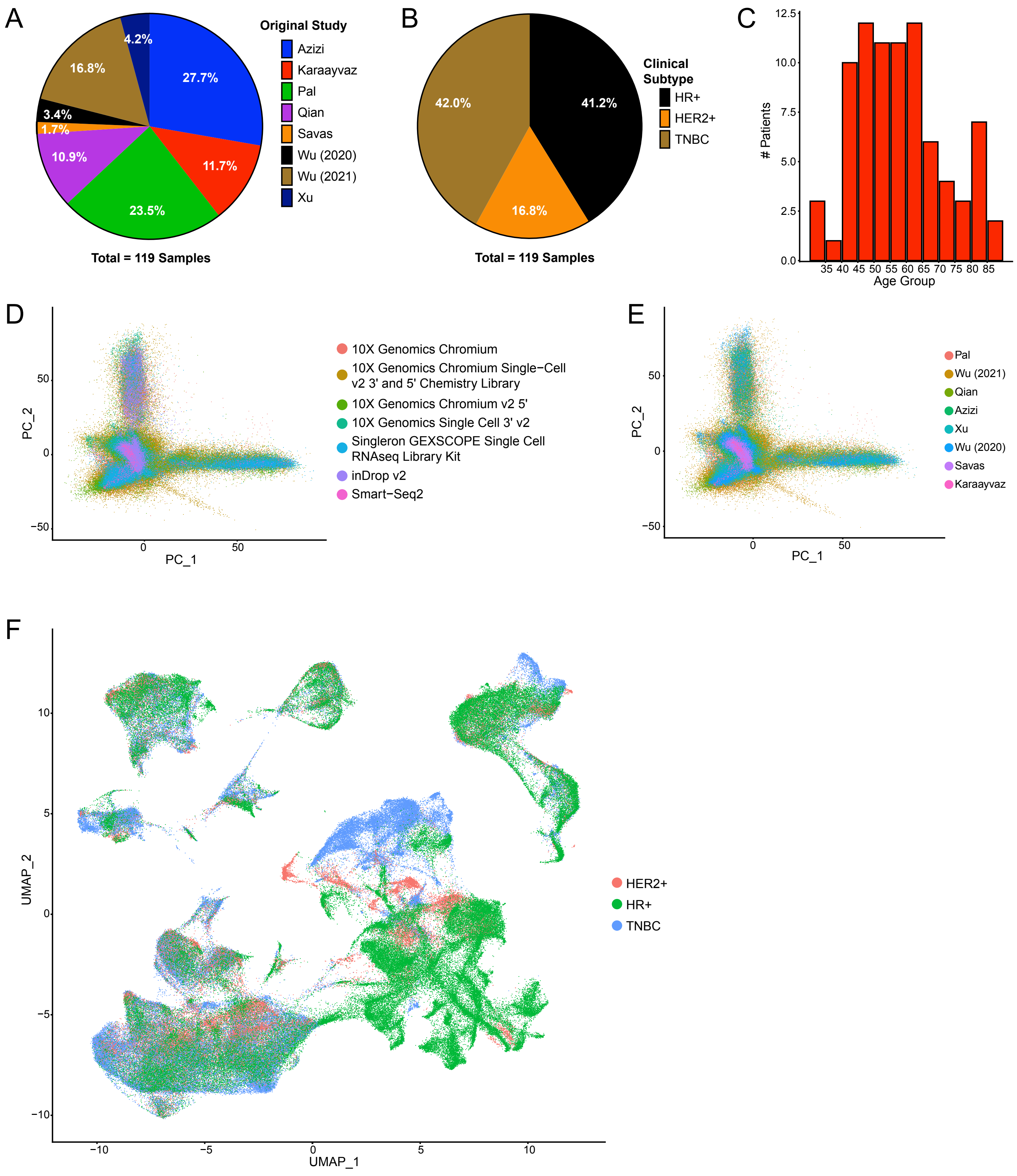

A
Original Study
Azizi
Karaayvaz
Pal
Qian
Savas
Wu (2020)
Wu (2021)
Xu
27.7%
11.7%
23.5%
10.9%
1.7%
3.4%
16.8%
4.2%
Total = 119 Samples
B
Clinical Subtype
HR+
HER2+
TNBC
41.2%
16.8%
42.0%
Total = 119 Samples
C
### Patients
Age Group
35 40 45 50 55 60 65 70 75 80 85
D
10X Genomics Chromium
10X Genomics Chromium Single−Cell v2 3' and 5' Chemistry Library
10X Genomics Chromium v2 5'
10X Genomics Single Cell 3' v2
Singleron GEXSCOPE Single Cell RNAseq Library Kit
inDrop v2
Smart−Seq2
PC_1
PC_2
E
Pal
Wu (2021)
Qian
Azizi
Xu
Wu (2020)
Savas
Karaayvaz
PC_1
PC_2
F
HER2+
HR+
TNBC
UMAP_1
UMAP_2

**Figure S1. Metadata and quality control of integrated scRNA-seq data**

A. Pie chart of composition of integrated scRNA-seq data by original study.
B. Pie chart of composition of integrated scRNA-seq data by clinical subtype. The proportion of clinical subtypes within this integrated dataset is close to real-life clinical subtype distributions.
C. Bar plot showing number of patients per age group. Most of the original datasets stayed within a sole age group, whereas the integrated dataset includes a much broader age range.
D. PCA plot of all cells colored by technology. No cluster is driven by a single technology, thus confirming there is no batch effect due to differing technologies.
E. PCA plot of all cells colored by the original study. No cluster is driven by a single study, thus confirming there is no batch effect due to different studies.
F. UMAP visualization of integrated scRNA-seq data grouped by clinical subtype. This shows lineage drives clustering of non-epithelial populations, while epithelial populations cluster by clinical subtype. This matches the observed subtype clustering seen in other datasets.

### Figure S2

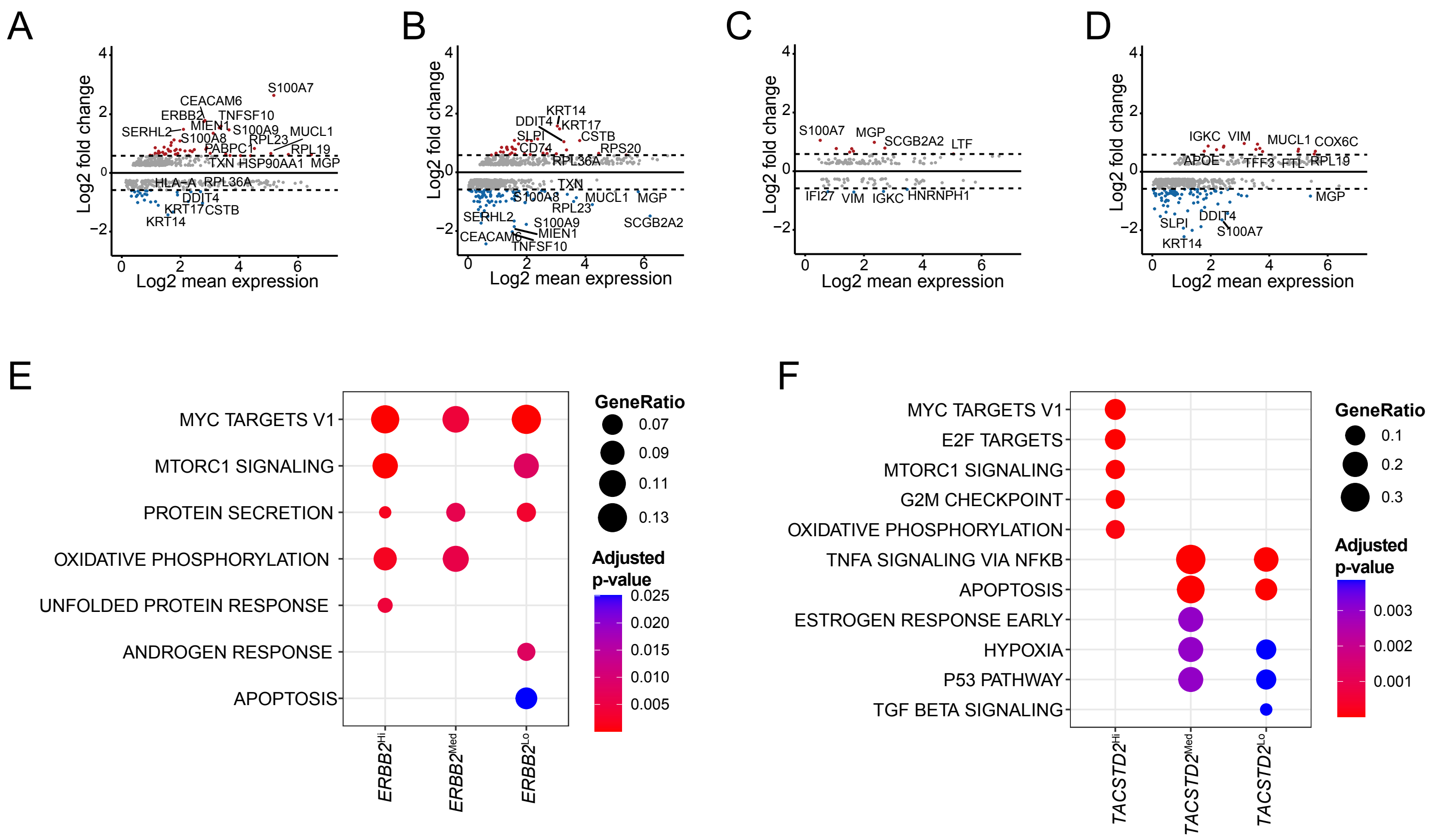
A
B
C
D
E
F
Log2 fold change
Log2 mean expression
S100A7
CEACAM6
ERBB2
TNFSF10
MIEN1
S100A9
MUCL1
SERHL2
S100A8
RPL23
PABPC1
RPL19
TXN
HSP90AA1
MGP
HLA-A
RPL36A
DDIT4
KRT17
CSTB
KRT14
SLPI
CD74
RPS20
SCGB2A2
LTF
IFI27
VIM
IGKC
HNRNPH1
COX6C
APOE
TFF3
FTL
MYC TARGETS V1
MTORC1 SIGNALING
PROTEIN SECRETION
OXIDATIVE PHOSPHORYLATION
UNFOLDED PROTEIN RESPONSE
ANDROGEN RESPONSE
APOPTOSIS
GeneRatio
Adjusted p-value
ERBB2Hi
ERBB2Med
ERBB2Lo
E2F TARGETS
G2M CHECKPOINT
TNFA SIGNALING VIA NFKB
ESTROGEN RESPONSE EARLY
HYPOXIA
P53 PATHWAY
TGF BETA SIGNALING
TACSTD2Hi
TACSTD2Med
TACSTD2Lo

**Figure S2. Differential gene expression and gene set enrichment analyses for each *ERBB2* and *TACSTD2* population.**

G. MA plot showing differentially expressed genes between *ERBB2*$^{Med}$ vs. *ERBB2*$^{Hi}$ and *ERBB2*$^{Lo}$ cells (Bonferroni adjusted p-value < 0.05).

H. MA plot showing differentially expressed genes between *ERBB2*$^{Lo}$ vs. *ERBB2*$^{Hi}$ and *ERBB2*$^{Med}$ cells (Bonferroni adjusted p-value < 0.05).

I. MA plot showing differentially expressed genes between *TACSTD2*$^{Med}$ vs. *TACSTD2*$^{Hi}$ and *TACSTD2*$^{Lo}$ cells (Bonferroni adjusted p-value < 0.05).

J. MA plot showing differentially expressed genes between *TACSTD2*$^{Lo}$ vs. *TACSTD2*$^{Hi}$ and *TACSTD2*$^{Med}$ cells (Bonferroni adjusted p-value < 0.05).

K. Gene set enrichment of the differentially expressed genes by *ERBB2*$^{Hi}$, *ERBB2*$^{Med}$, and *ERBB2*$^{Lo}$ cells. Significantly enriched gene sets from the MSigDB HALLMARK collection are shown (Benjamini-Hochberg adjusted p-value < 0.05).

L. Gene set enrichment of the differentially expressed genes by *TACSTD2*$^{Hi}$, *TACSTD2*$^{Med}$, and *TACSTD2*$^{Lo}$ cells. Significantly enriched gene sets from the MSigDB HALLMARK collection are shown (Benjamini-Hochberg adjusted p-value < 0.05).

### Figure S3

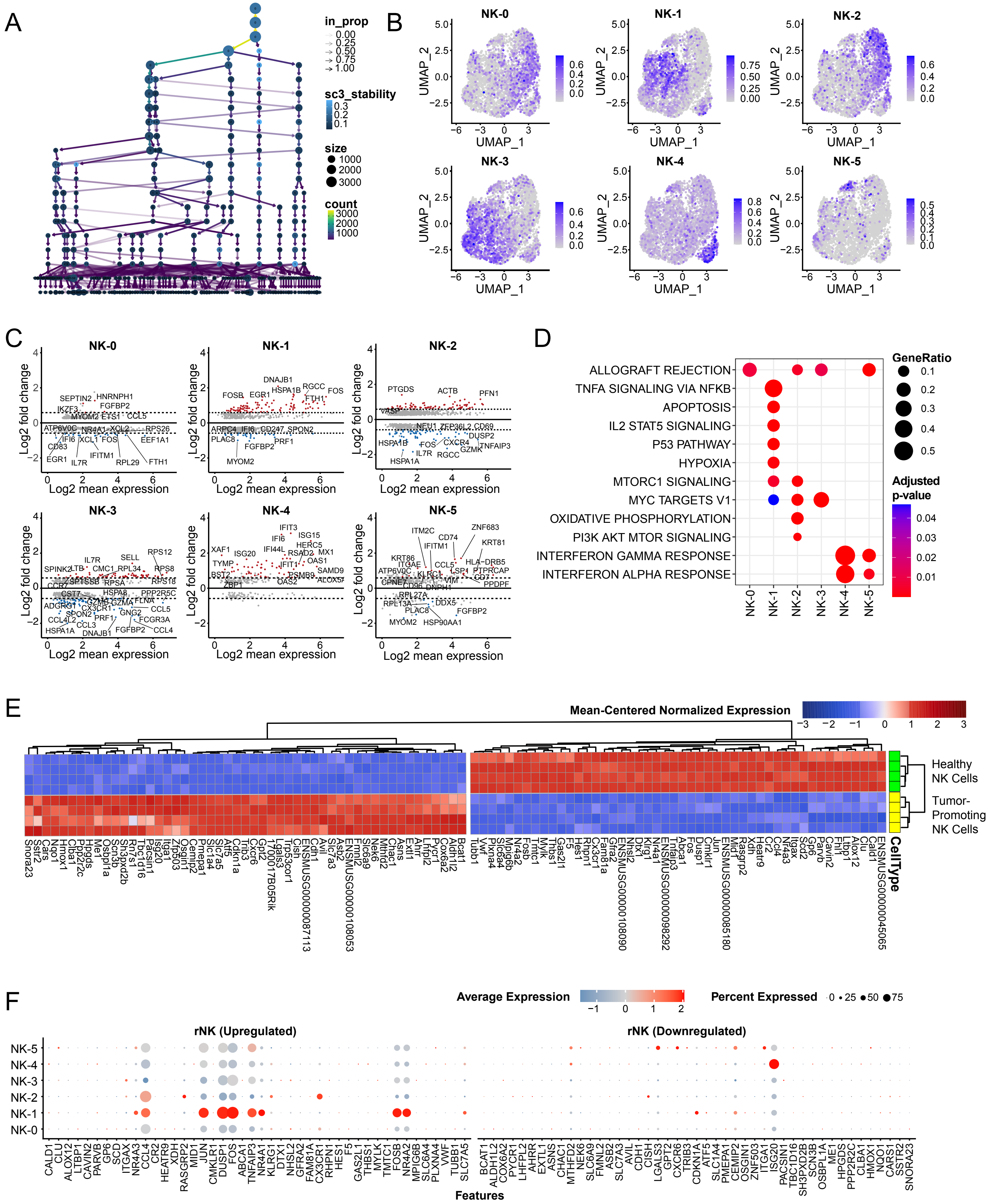
A
in_prop
0.00
0.25
0.50
0.75
1.00
sc3_stability
0.3
0.2
0.1
size
1000
2000
3000
count
3000
2000
1000
B
NK-0
NK-1
NK-2
NK-3
NK-4
NK-5
UMAP_1
UMAP_2
C
Log2 fold change
Log2 mean expression
D
ALLOGRAFT REJECTION
TNFA SIGNALING VIA NFKB
APOPTOSIS
IL2 STAT5 SIGNALING
P53 PATHWAY
HYPOXIA
MTORC1 SIGNALING
MYC TARGETS V1
OXIDATIVE PHOSPHORYLATION
PI3K AKT MTOR SIGNALING
INTERFERON GAMMA RESPONSE
INTERFERON ALPHA RESPONSE
GeneRatio
0.1
0.2
0.3
0.4
0.5
Adjusted p-value
0.04
0.03
0.02
0.01
E
Mean-Centered Normalized Expression
Healthy NK Cells
Tumor-Promoting NK Cells
CellType
F
Average Expression
Percent Expressed
0
25
50
75
rNK (Upregulated)
rNK (Downregulated)
Features

**Figure S3. Unsupervised clustering of NK cell subsets and rNK signature development.**

A. Clustree diagram showing unsupervised clustering of all cancer epithelial cells across 15 resolutions (0.01, 0.05, 0.08, 0.1, 0.2, 0.3, 0.4, 0.5, 0.6, 0.7, 1.0, 1.3, 1.6, 1.8, 2.0).
B. Feature plots showing expression of NK subset markers across all NK cells in our integrated dataset.
C. MA plots showing differentially expressed genes between individual NK cell subsets and all other NK cell subset types (Bonferroni adjusted p-value $< 0.05$).
D. Gene set enrichment of the differentially expressed genes by each NK cell subset. Significantly enriched gene sets from the MSigDB HALLMARK collection are shown (Benjamini-Hochberg adjusted p-value $< 0.05$).
E. Heatmap showing z-scores for the variance-stabilized transformed expression of differentially expressed genes between healthy NK cells and tumor-promoting NK cells from previous study.
F. Bubble heatmap showing expression of upregulated and downregulated human rNK orthologs for each major NK cell subset.

### Figure S4

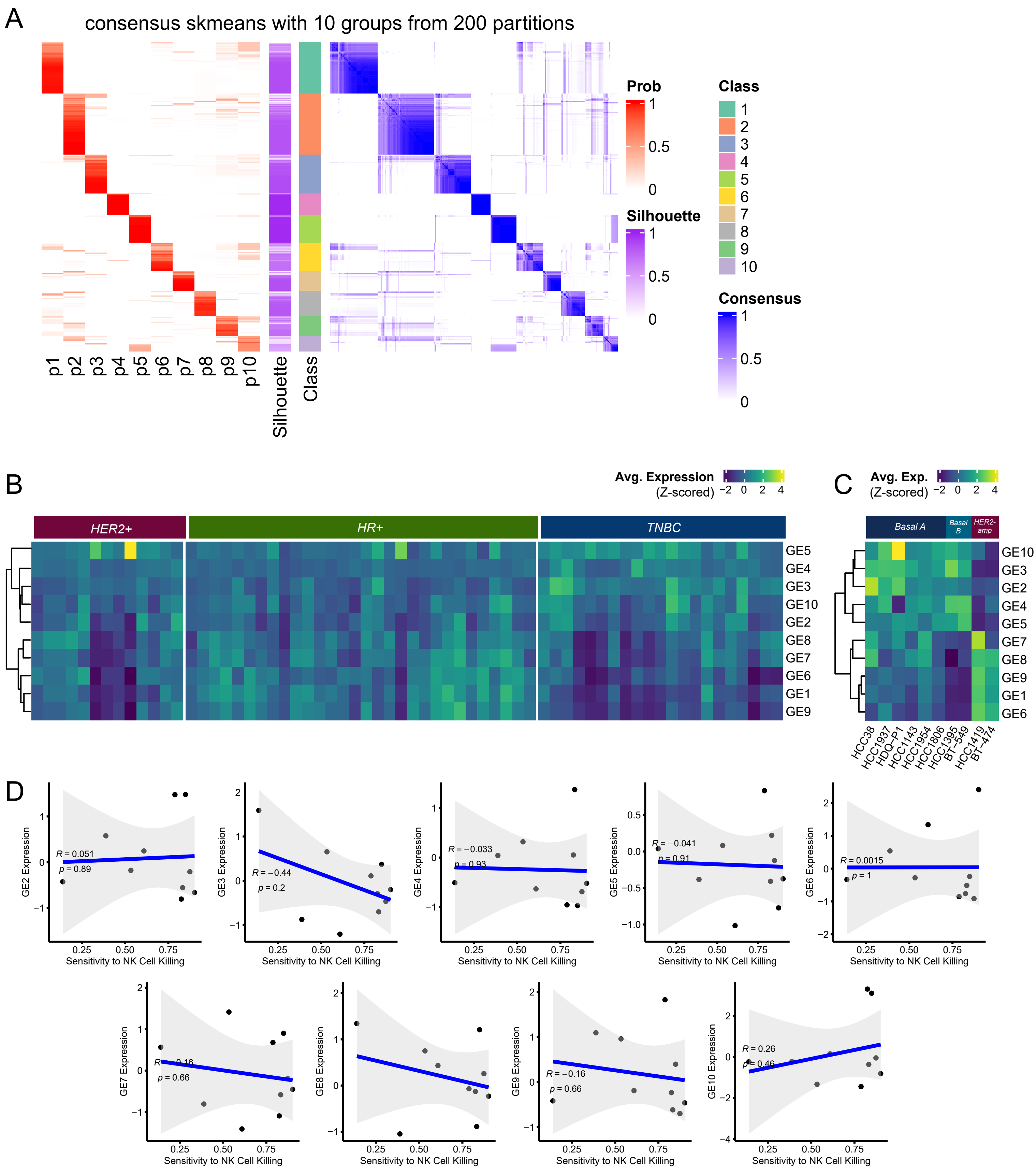
A
consensus skmeans with 10 groups from 200 partitions
p1
p2
p3
p4
p5
p6
p7
p8
p9
p10
Silhouette
Class
Prob
1
0.5
0
Silhouette
1
0.5
0
Class
1
2
3
4
5
6
7
8
9
10
Consensus
1
0.5
0
B
Avg. Expression
(Z-scored)
−2 0 2 4
HER2+
HR+
TNBC
GE5
GE4
GE3
GE10
GE2
GE8
GE7
GE6
GE1
GE9
C
Avg. Exp.
(Z-scored)
−2 0 2 4
Basal A
Basal B
HER2-amp
GE10
GE3
GE2
GE4
GE5
GE7
GE8
GE9
GE1
GE6
HCC38
HCC1937
HDQ−P1
HCC1143
HCC1954
HCC1806
HCC1395
BT−549
HCC1419
BT−474
D
GE2 Expression
R = 0.051
p = 0.89
GE3 Expression
R = −0.44
p = 0.2
GE4 Expression
R = −0.033
p = 0.93
GE5 Expression
R = −0.041
p = 0.91
GE6 Expression
R = 0.0015
p = 1
GE7 Expression
R = −0.16
p = 0.66
GE8 Expression
GE9 Expression
R = −0.16
p = 0.66
GE10 Expression
R = 0.26
p = 0.46
Sensitivity to NK Cell Killing
0.25
0.50
0.75

**Figure S4. Generation and characterization of the 10 gene elements of cancer epithelial cell heterogeneity.**

A. Spherical k-means (skmeans) consensus clustering of the Jaccard similarities between signatures of cancer epithelial cell ITTH, showing the probability (p1-p10) of each generated signature of being assigned to one of 10 classes. Silhouette scores are shown for each class or GE.
B. Heatmap of average z-scored expression of each of the 10 GEs across cancer epithelial cells in each sample in our integrated dataset.
C. Heatmap of average z-scored expression of each of the 10 GEs across 10 cell lines derived from primary breast tumors. Cell lines are annotated by molecular subtype (basal A, basal B, HER2-amplified).
D. Scatterplots showing Pearson correlations of expression of GEs with limited predicted interactions with NK cells (all but GE1) and sensitivity to NK cell killing across 10 cell lines derived from primary breast tumors (Benjamini-Hochberg adjusted p-values > 0.05).

### Figure S5

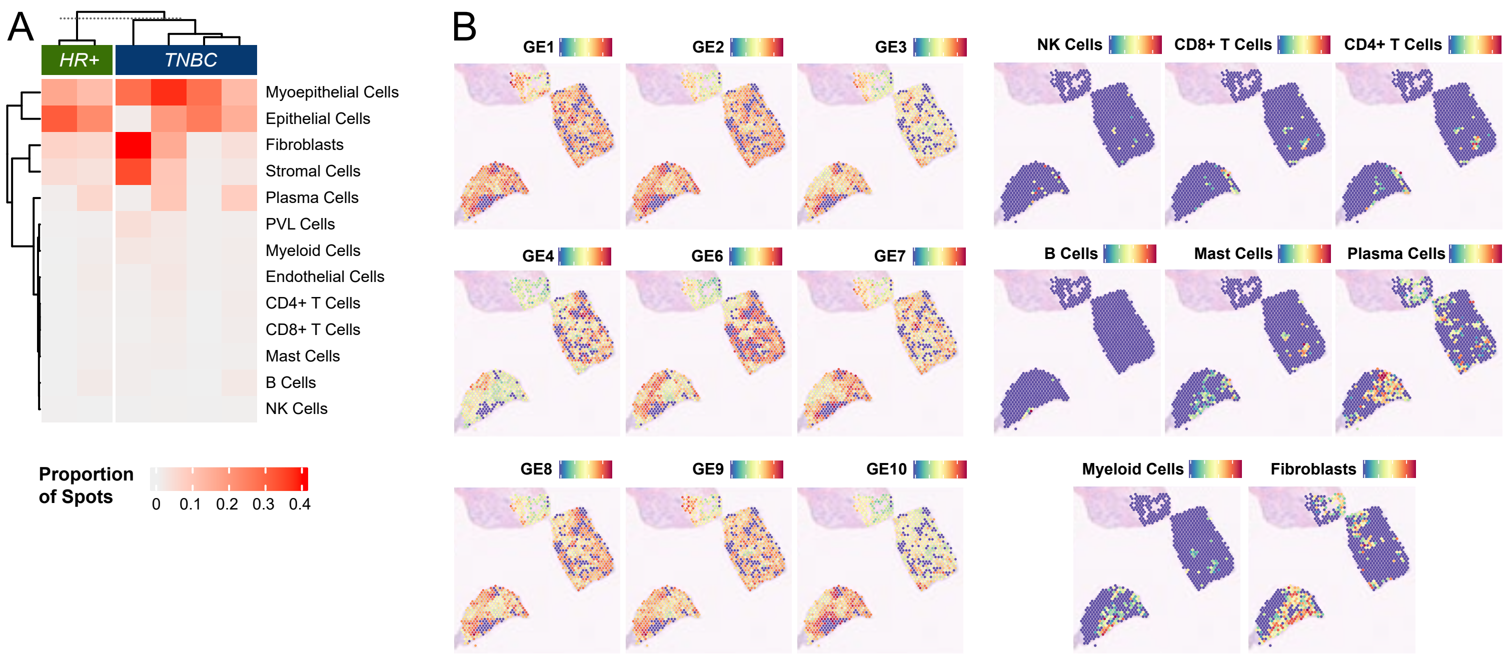
A
HR+
TNBC
Myoepithelial Cells
Epithelial Cells
Fibroblasts
Stromal Cells
Plasma Cells
PVL Cells
Myeloid Cells
Endothelial Cells
CD4+ T Cells
CD8+ T Cells
Mast Cells
B Cells
NK Cells
Proportion of Spots
0
0.1
0.2
0.3
0.4
B
GE1
GE2
GE3
NK Cells
CD8+ T Cells
CD4+ T Cells
GE4
GE6
GE7
B Cells
Mast Cells
Plasma Cells
GE8
GE9
GE10
Myeloid Cells
Fibroblasts

**Figure S5. Deconvolution of spatial transcriptomics samples and spatial resolution of the 10 GEs and GE-immune interactions.**

A. Heatmap showing the proportion of spatial tumor sample spots within a sample that contain each of the GEs and immune or stromal cell populations.
B. For a representative TNBC sample, UCell signature scores of each GE overlaid onto spatial tumor sample spots with >10% presence of cancer epithelial cells. UCell signature scores of immune and stromal populations overlaid onto respective spots annotated by integration with the scRNA-seq dataset.
